## Supplemental Figures 1-6 for "Ectoderm to mesoderm transition by downregulation of actomyosin contractility"

### Figure S1

A

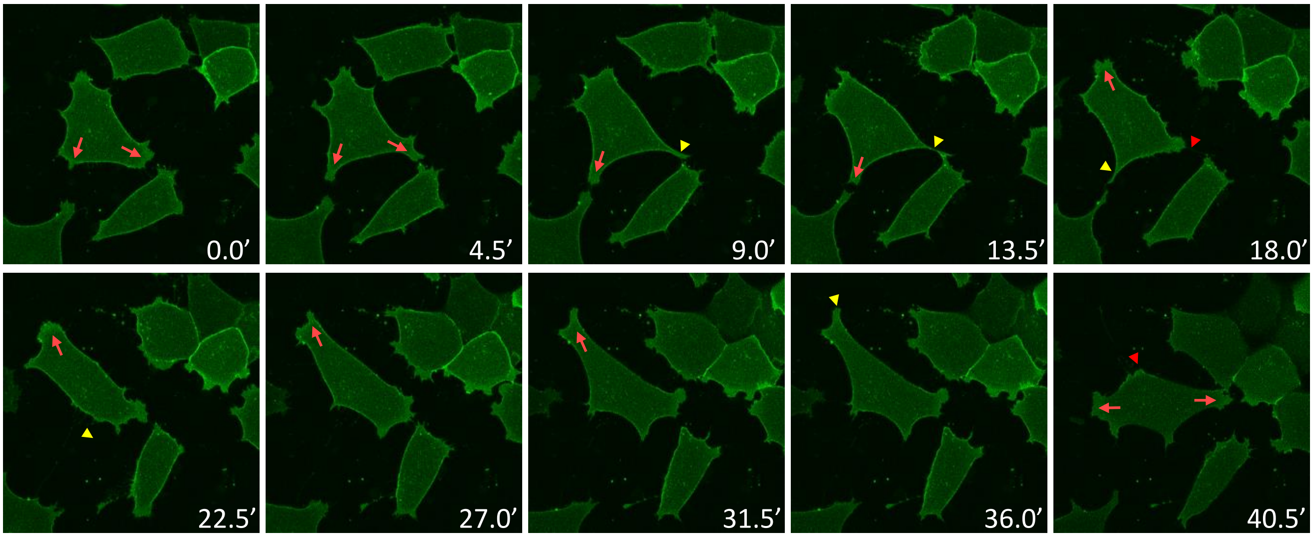


**A) Mode of mesoderm locomotion** (Related to Figure 1) Consecutive frames from time lapse of mYFP labelled mesoderm cells migrating on FN. The behaviour of the central cell is highlighted: The cell emits one or multiple protrusions (red arrows). One of the protrusions becomes a tail (yellow arrowhead) as the cell stretches toward another direction, and eventually retracts (red arrowheads).

B

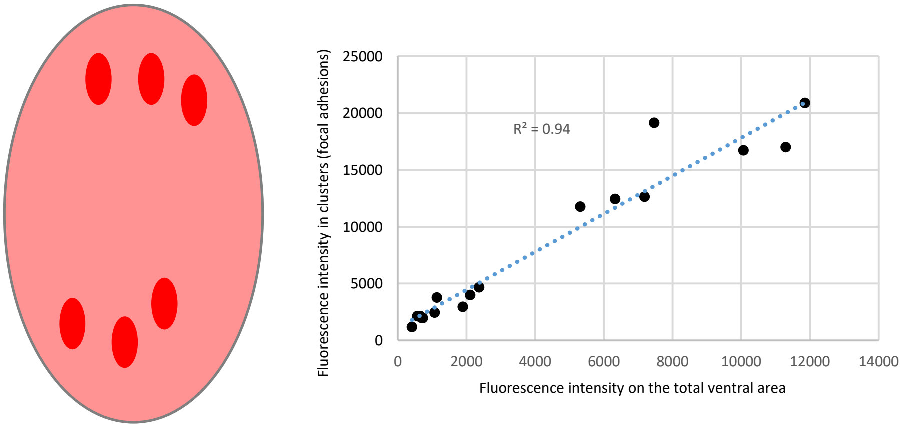


**B) Quantification of accumulation of Vinculin-Cherry in focal adhesions: Linearity between fluorescence levels in focal adhesion and total intensity** (Related to Figure 1A-D) Because Vinculin-Cherry expression levels vary from cell to cell, quantification was performed for individual cells by measuring fluorescence in bright clusters (corresponding to focal adhesions) and in the total ventral cell surface (pink on the diagram). The plot shows the average intensity of the ventral surface versus the average intensity in focal adhesions for control mesoderm cells in one experiment, each dot corresponding to a single cell. It shows that accumulation at focal adhesions is proportional to total expression levels over a wide range. Linearity was similarly verified for each experiment.

### Figure S2

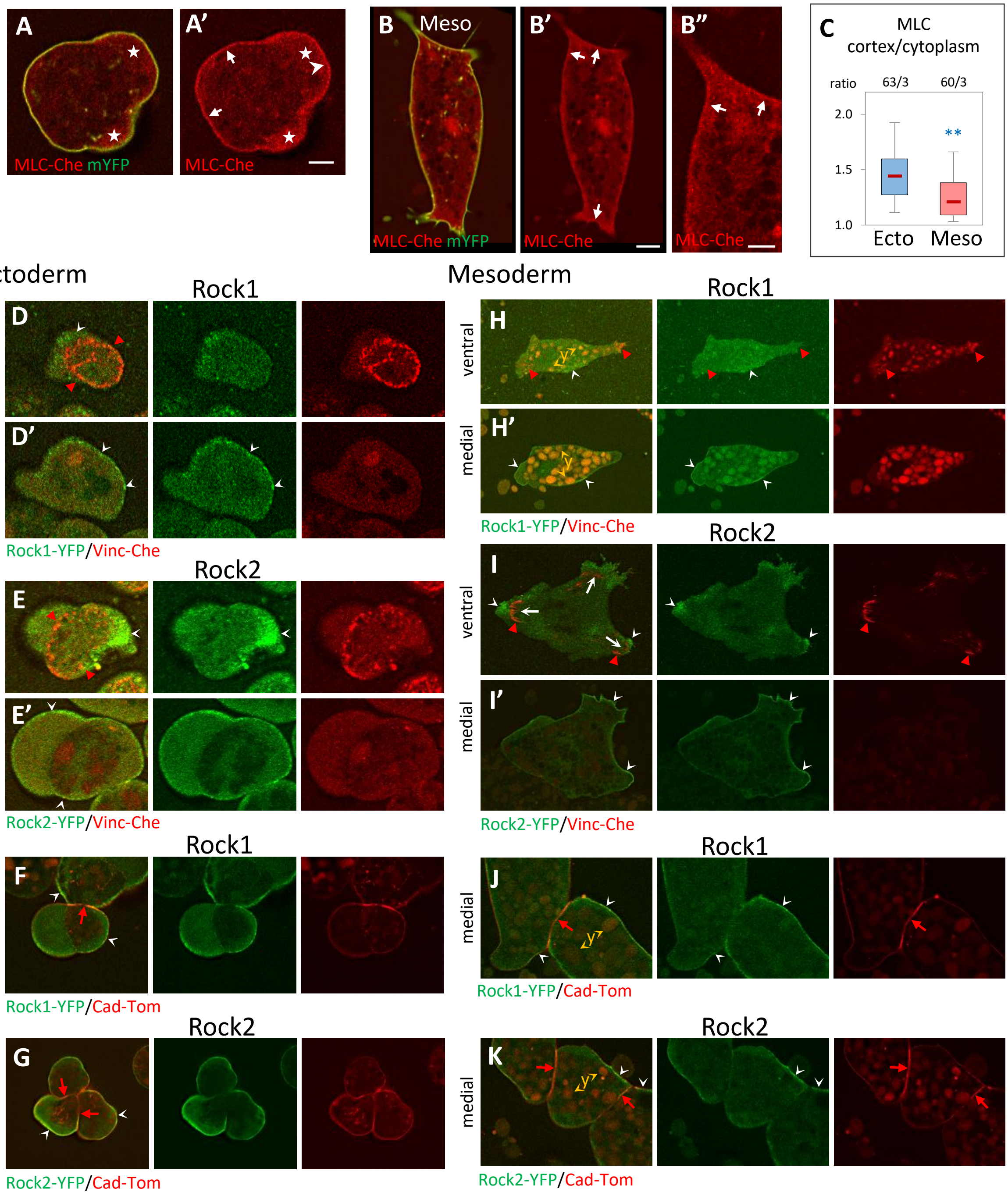


**Localization of MLC and Rock** (Related to Figure 2)

**A-C) Differential MLC accumulation at the cell cortex.** Ectoderm and mesoderm cells expressing MLC-Cherry (MLC-Che) and mYFP. A) Ectoderm cells show strong accumulation around the cell body (arrows) and part of the blebs (arrowhead). B) Mesoderm cells show irregular cortical MLC, mostly at the concave regions near or between protrusion. C) Quantification of cortical MLC, expressed as the ratio of cortical /cytoplasmic fluorescence intensities. Blebs and protrusions were excluded from the measurements. Statistical comparison using two-sided Student's t-test. Scale bars: A' 5μm, B' 10μm, B" 5μm.

**D-K) Subcellular localization of Rock1-YFP and Rock2-YFP in ectoderm and mesoderm cells.** Selected single planes from live confocal microscopy, either near the glass (ventral), or about 5-10μm above (medial). Concave white arrowheads point at examples of Rock1/2 accumulation. D,E,H,I) Localization relative to the cell cortex and to vinculin-Cherry labelled cell-matrix adhesive structures (red arrowheads). F,G,J,K) Localization relative to cell-cell contacts, marked by cadherin-dTomato (red arrows). D,E) In the ectoderm, Rock1 and 2 have both a cortical localization. Levels are low on the ventral side inside the adhesive ring, but stronger outside of the ring, particularly for Rock2. F,G) Levels are very low at cell-cell contacts. H,I) In the ventral face of mesoderm cells, Rock1 tend to be enriched in the central part, Rock2 at the periphery of the protrusions. Both are low at FAs. They both accumulate at the cortex along cell free edges (medial planes). J,K) Levels are low at cell-cell contacts. Y: autofluorescence of yolk platelets, abundant in mesoderm cells.

### Figure S3

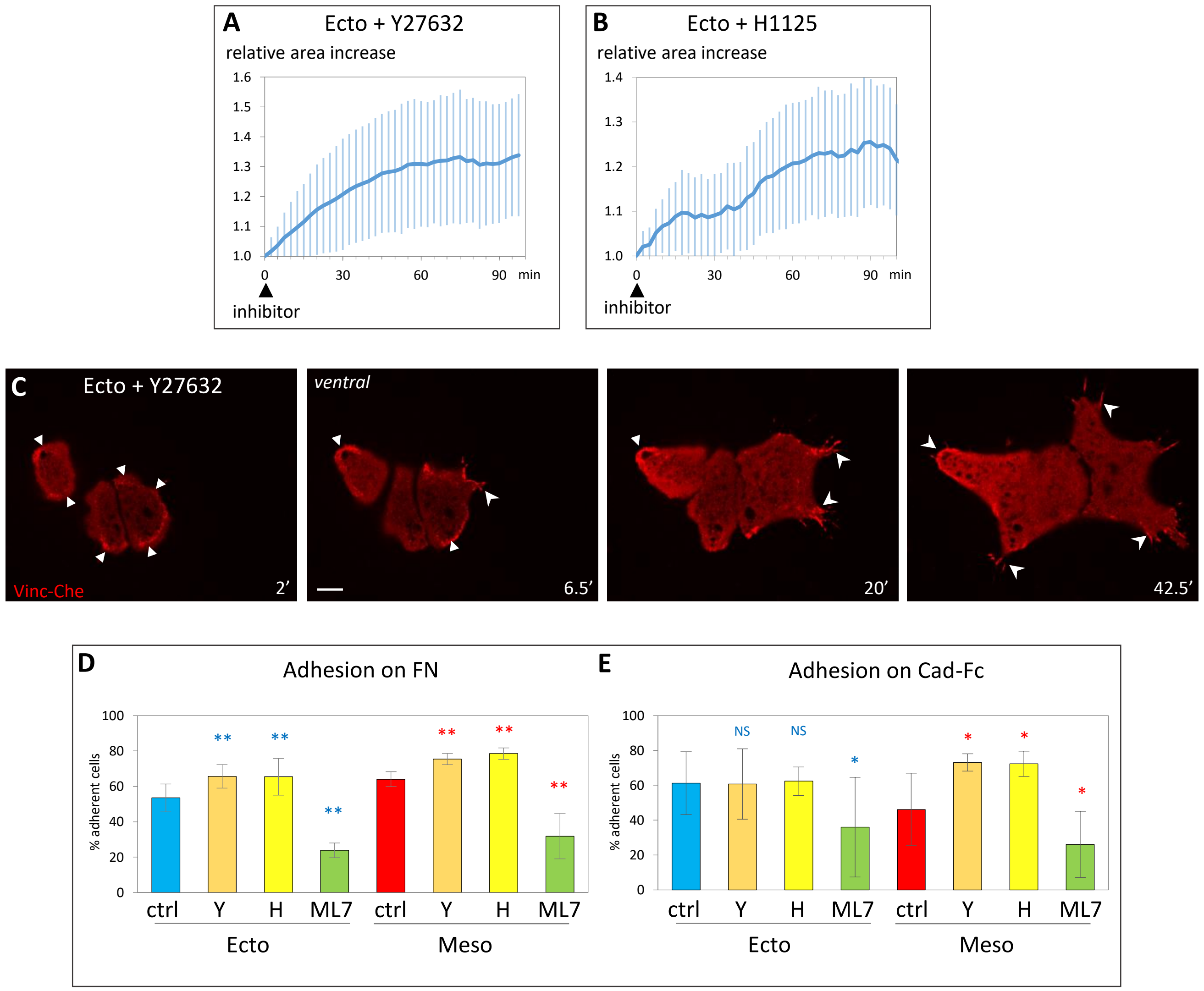


Related to Figure 2: **A,B) Area expansion** for single cells after treatment with Rock inhibitors Y27632 (50µM) and H1125 (1µM). Average and SD of 107 cells (A) and 34 cells (B). **C) Changes in vinculin distribution.** Images from a time lapse movie of a small group of three cells expressing Vinculin-Cherry, treated at time = 0 with Y27632. Filled arrowheads: ring-like adhesion; Concave arrowheads: FAs. Scale bars: 10µm. **D,E) Opposite effects of Rock and MLCK inhibition on cell adhesion.** Ectoderm and mesoderm adhesion to FN or cadherin was measured after treatment with Rock inhibitors Y27632 (Y, 50µM), H1125 (H, 1µM), or the MLCK inhibitor ML7. 5 experiments, total 1000-2000 cells/conditions. Statistical comparison to control ectoderm or mesoderm, comparing the % adherent cells/experiment, pairwise two-sided Student's t-test.

### Figure S4

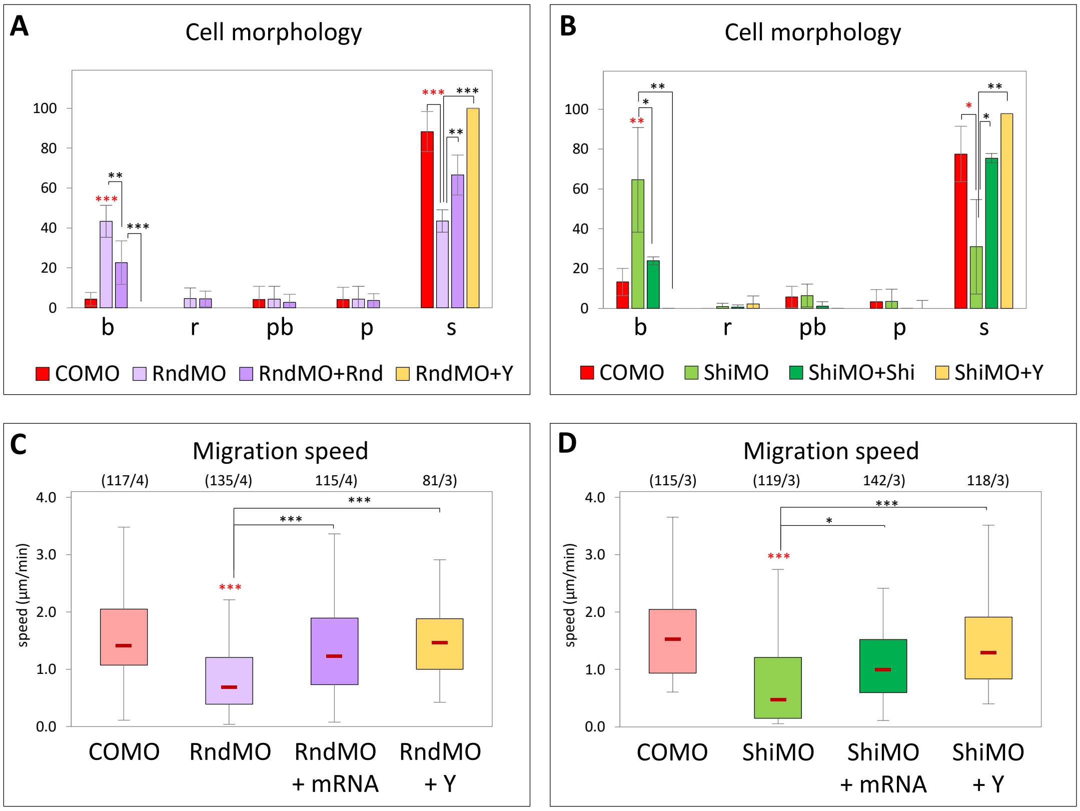


**A-D) Rescue of Rnd1MO and ShiMO spreading and migration phenotypes** (Related to Figure 4)**.** 4-cell stage embryos were injected in the dorsal side with COMO, RndMO, RndMO + YFP-Rnd1 mRNA (rescue), ShiMO, or ShiMO + YFP-Shirin mRNA (rescue). Dissociated mesoderm cells were plated on FN and time lapse movies were recorded. The fourth condition represents RndMO or ShiMO cells treated with 50µM Y27632 Rock inhibitor (Y). Statistical comparions: One-way ANOVA followed by Tukey's HSD post hoc test. Red asterisks: Comparison to COMO.

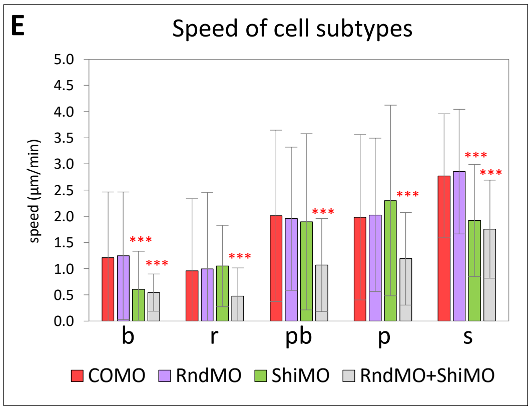


**E) Migration speed for different cell morphology categories** (Related to Figure 4). Analysis of data from figure 3K. Red asterisks: Comparison to COMO. one-way ANOVA followed by Tukey's HSD post hoc test.

### Figure S5

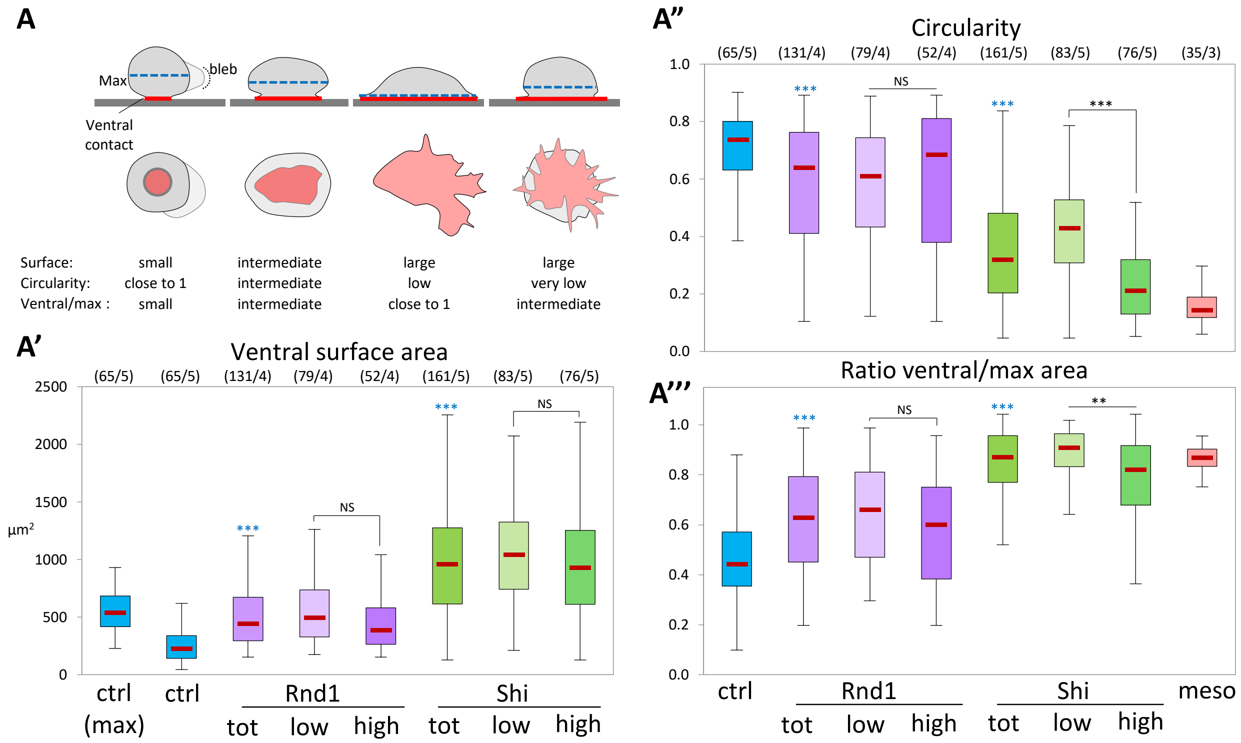


**A) Morphometry of Rnd1 and Shirin induced spreading of ectoderm cells** (Related to Figure 4).

The diagrams illustrate typical cell shapes. Corresponding images can be found in main Figure 5A-E. These shapes were analysed based on the following parameters: A') Area of the ventral contact surface (red in the schemes in A). A'') Circularity of the ventral surface, which depends both on the roundness and regularity/convolution of the shape. A''') Ratio between the ventral area and the maximal cell area, calculated from maximal z projections. Blebs were excluded from measurements. Rnd1 and Shirin-expressing cells were here subdivided in two categorises, low and high-expression, based on the YFP fluorescence intensity. Note that these two categories overlap but are not equivalent Rnd1 expression levels had no significant impact on any parameter. Shirin expression had no effect on contact surface area, but high levels stimulated formation of convoluted protrusions (lower circularity) but decreased ventral/max area, reflecting the fact that many of them rounded up (4th cell shape in panel A, see main figure 5E).

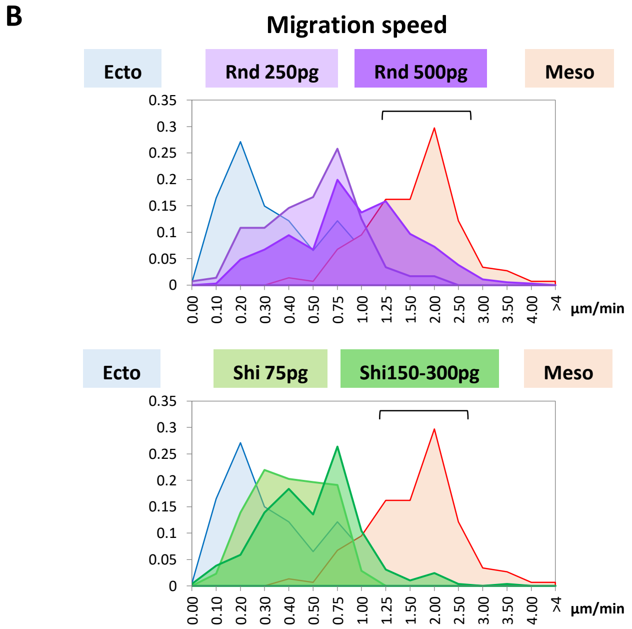


B) Histograms showing distribution of speed for ectoderm cells expressing Rnd1 or Shirin, compared to wild type ectoderm and mesoderm (Related to Figure 4). Brackets: Range of high speed, comparable to mesoderm, achieved mainly by Rnd1-expressing cells.

### Figure S6

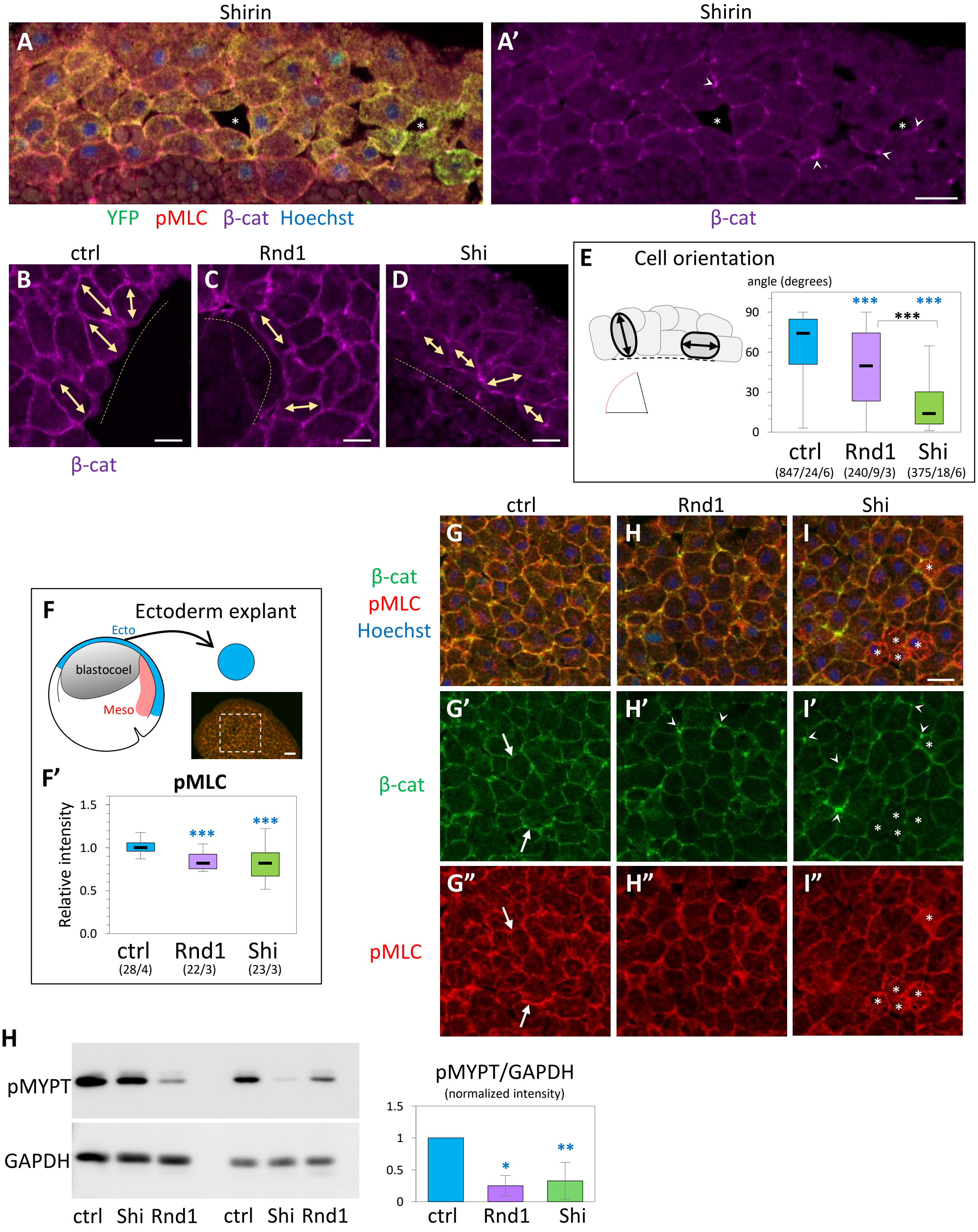


**Effect of Rnd1 and Shirin ectopic expression** (Related to Figure 6). **A) Loosening of ectoderm tissue upon expression of Shirin.** Immunostained section of a YFP-Shirin expressing embryo showing a loosely organized ectoderm, characterized by the presence of large intercellular spaces (asterisks) and heterogenous β-catenin signal, weak signal along membranes except for strong local concentrations (arrowheads). Scale bar, 10µm. **B-E) Cell orientation**. B-D) The main axis of deep ectoderm cells (double arrows) tend to orient roughly perpendicular to the inner surface of the tissue (dashed line). Rnd1-expressing cells show variable orientation. Shirin-expressing cells align parallel to the surface. Scale bars, 10µm. E) Quantification of the angle between the cell axis and the tissue interface. Numbers into brackets correspond to number of cells/embryos/experiments.

**F-I) Analysis of β-catenin (green) and pMLC (red) in ectoderm explants**. F) Diagram, section of an control ectoderm explant (scale bar, 50µm) and quantification. Statistical comparison using one-way ANOVA followed by Tukey's HSD post hoc test. G-I) Examples of ectoderm explants. G) β-catenin and pMLC signal along cell edges is highest in control (arrows). H,I) Explants expressing Rnd1 or Shirin. β-catenin tends to accumulate at cell vertices (concave arrowheads). pMLC levels are lower except for some cells (I, asterisks) that have rounded up, and display high pMLC throughout the cell. Little to no β-catenin is seen between the round cells. Scale bars, 20µm. **H) Effect of Rnd1 and Shirin expression on phosphorylation of MYPT.** Dissected ectoderm tissues were analysed by Western Blot. GAPDH was used as loading control, and the pMYPT signal was expressed as relative ratio, normalized to ectoderm control set to 1.0. Three independent experiments, statistical analysis using one sample, two-sided *t-test*.

### Figure S7

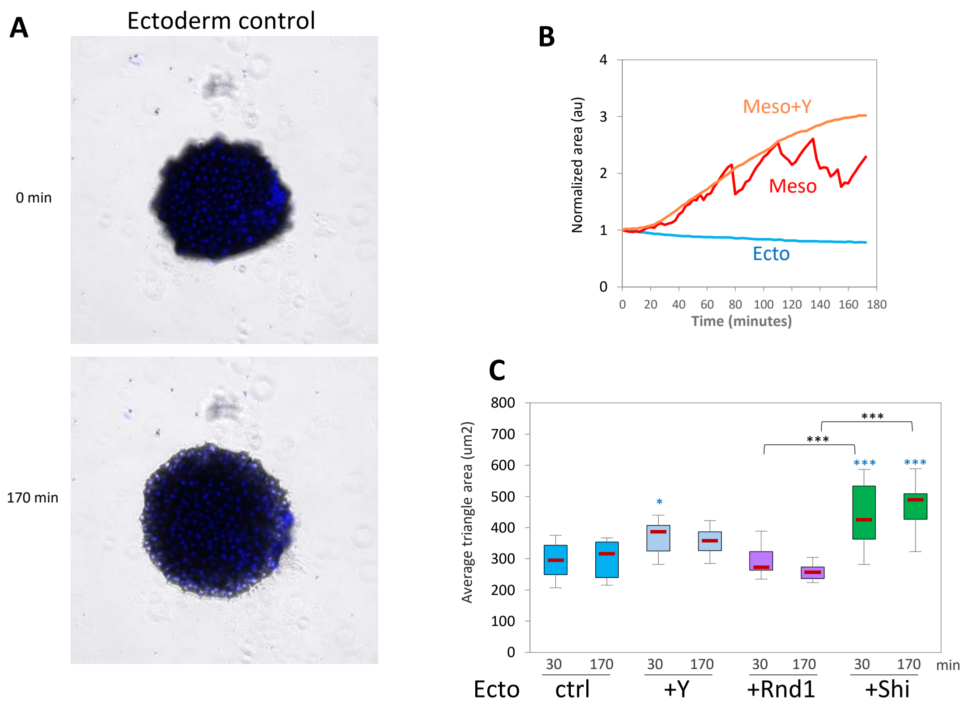


**Figure S7** (Related to Figure 9): A) Example of ectoderm explant showing late partial spreading, which is only observed beyond the 120min. B) Examples of traces for single explants, illustrating the irregular expansion of mesoderm explants interrupted by retractions. In contrast, expansion of Y26862- treated mesoderm is smooth. C) Quantification of average triangle size at the initiation of spreading (30 min) and the end of the time lapse (170 min).

**Figure S8 (next page). Knockdown of Rnd1 or Shirin affect collective properties of mesoderm tissue explants** (Related to Figure 9). Analysis of spreading, dispersion, and intercalation of mesodermal explants under various conditions was performed as for experiments presented in Figure 9. A-D) Control mesoderm, mesoderm treated with Y26862, and mesoderm from embryos injected with Rnd1 MO or Shirin MO. Red arrowheads in A and D indicate areas of large scale retractions (compare 85 and 170min). Scale bar: 100µm. E) Average time course curves with SD for the various experimental conditions. E') Corresponding relative spreading after 60min and 170min. F-K) Delaunay triangulation of nuclei in order to measure cell dispersion. F-I) Representative plots of triangulated nuclei after 170 minutes of imaging. X and Y labels mark the coordinates in µm, the colour coded scale bar indicates the area of the triangles in $\mu m^2$. J) Quantification of average triangle size at the initiation of spreading (30 min) and the end of the time lapse. K) Quantification of the relative change in triangle size over time calculated by dividing the average triangle area at 170 minutes by that at 30 minutes. L) Quantification of intercalation calculated by dividing number of nuclei at the ventral surface at 170 minutes by the number at 30 minutes. Statistical comparisons: One-way ANOVA followed by Tukey's HSD post hoc test.

### Figure S8

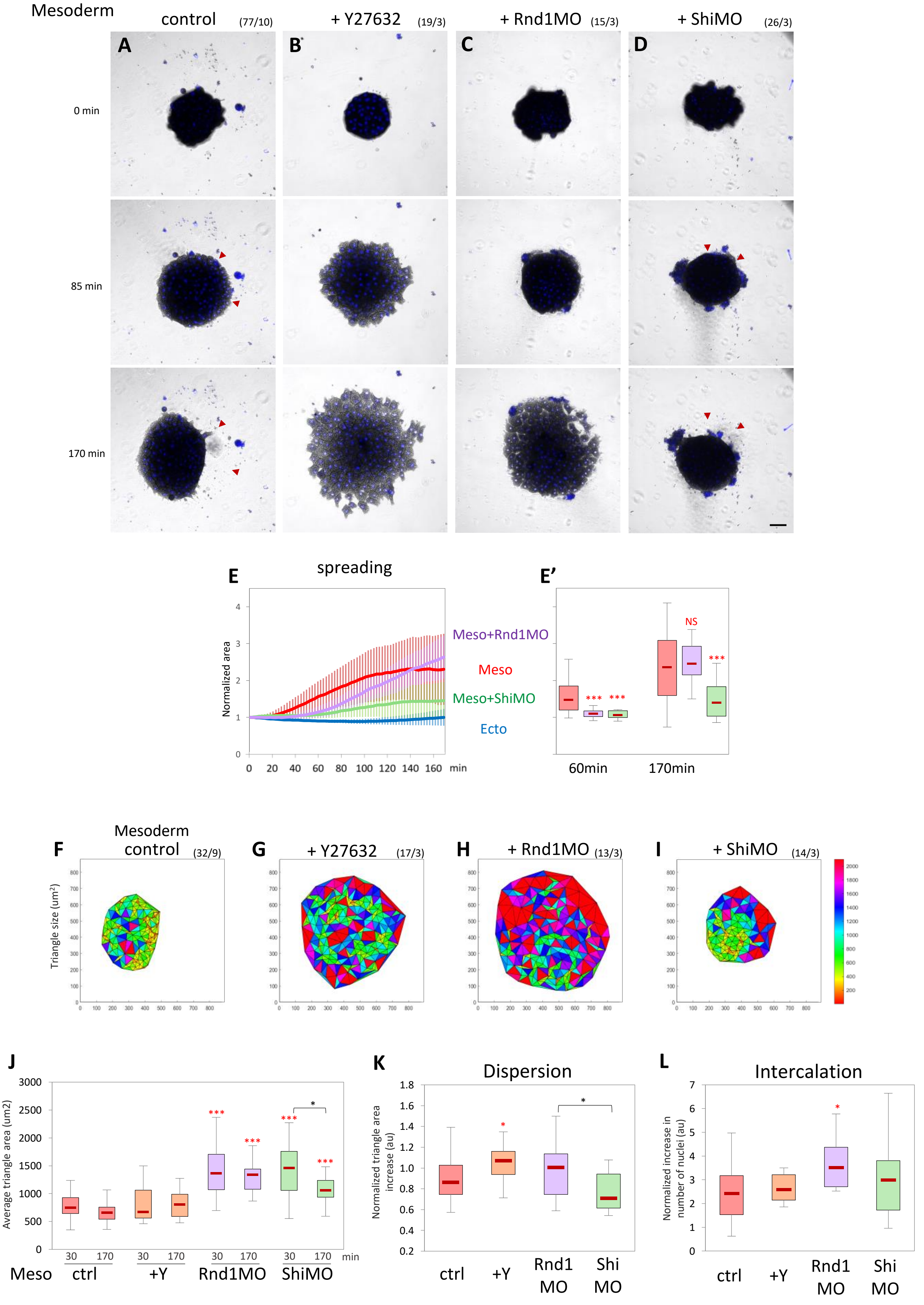

Mesoderm
control (77/10)
+ Y27632 (19/3)
+ Rnd1MO (15/3)
+ ShiMO (26/3)
A
B
C
D
0 min
85 min
170 min
E
spreading
Normalized area
Meso+Rnd1MO
Meso
Meso+ShiMO
Ecto
0 20 40 60 80 100 120 140 160 min
E'
NS
***
60min
170min
F
Mesoderm
control (32/9)
G
+ Y27632 (17/3)
H
+ Rnd1MO (13/3)
I
+ ShiMO (14/3)
Triangle size (um²)
J
Average triangle area (um2)
30 170 30 170 30 170 30 170 min
Meso ctrl +Y Rnd1MO ShiMO
K
Dispersion
Normalized triangle area increase (au)
ctrl +Y Rnd1 MO Shi MO
L
Intercalation
Normalized increase in number of nuclei (au)
ctrl +Y Rnd1 MO Shi MO
